# FateLimit quantifies the prediction horizon of cell fate

**DOI:** 10.64898/2026.06.22.733672

**Authors:** Ji-Yong Sung, Jae-Ho Cheong

## Abstract

Single-cell technologies have enabled increasingly detailed reconstruction of developmental trajectories, yet a fundamental question remains unresolved: when does future cellular identity become predictable from a cell’s current molecular state? Existing approaches infer lineage relationships, transition probabilities or future transcriptional dynamics, but do not directly quantify the emergence of fate predictability during cellular state transitions. Here we present FateLimit, an information-theoretic framework for measuring the temporal dynamics of cell-fate predictability from single-cell omics data. FateLimit combines probabilistic fate assignment, fate entropy and mutual information to quantify how information about future cellular outcomes is encoded in present molecular states. We introduce two quantitative descriptors: the Fate Information Half-Life (FIHL), which measures the characteristic timescale of fate-information dynamics, and the Prediction Horizon (PH), defined as the earliest developmental stage at which observed fate predictability exceeds the 95th percentile of a permutation-derived null distribution. We applied FateLimit across developmental, lineage-tracing and reprogramming systems, including pancreatic endocrinogenesis, CellTag reprogramming, human hematopoiesis and zebrafish embryogenesis. Across all datasets, FateLimit identified significant fate information and reproducible prediction horizons that were robust to cell-state representation, lineage structure and biological context. Comparative analysis revealed that prediction horizons differ substantially among cellular lineages, indicating that distinct developmental programs acquire predictive information at different rates. FateLimit establishes a general framework for quantifying the predictability of future cellular identity from present molecular states. By transforming developmental trajectories into predictability landscapes, FateLimit enables systematic comparison of commitment dynamics across biological systems and establishes prediction horizons as a quantitative measure of cell-fate determination.

## Introduction

Cell fate determination is a fundamental process underlying development, tissue homeostasis, regeneration and disease progression. ^1,2^ Recent advances in single-cell omics technologies have enabled unprecedented characterization of cellular states and transitions at high resolution. ^3,4^ In parallel, computational approaches such as RNA velocity, ^5^ trajectory inference, lineage tracing integration and fate mapping algorithms have substantially improved our ability to predict future cellular states from molecular measurements.^6,7^ These methods have transformed single-cell biology from a descriptive discipline into a predictive science. Despite these advances, a fundamental question remains unresolved: to what extent is cell fate predictable? Existing approaches are primarily designed to maximize predictive accuracy or reconstruct developmental trajectories. As a result, prediction performance is often interpreted as a proxy for biological determinism. ^8,9^

However, limited prediction accuracy may arise from multiple sources, including insufficient measurements, incomplete molecular characterization, model limitations or intrinsic biological variability. Consequently, current frameworks cannot distinguish whether an observed prediction error reflects technical limitations or a fundamental limit of predictability in cell fate determination.^10^ This challenge is increasingly important as single-cell technologies evolve toward multiomic measurements. The integration of transcriptomic, epigenomic, proteomic and lineage information is expected to improve prediction accuracy.^11^ Nevertheless, it remains unknown whether progressively richer molecular measurements can eventually achieve complete predictability of cellular outcomes, or whether a residual uncertainty persists even after incorporating all available information.^10^ Addressing this question requires a conceptual shift from predicting cell fate to quantifying the limits of cell fate prediction Here we introduce FateLimit, a computational framework for quantifying the prediction horizon of cell fate from single-cell data. Rather than focusing on the development of increasingly accurate predictive models, FateLimit estimates how far into the future cellular outcomes remain predictable and quantifies the information about future fate contained within a present cellular state. ^12^

We define three complementary quantities: fate entropy, which measures uncertainty in future cellular outcomes; Fate Information Half-Life (FIHL), which measures the characteristic timescale over which fate information changes during cellular state transitions; and Prediction Horizon (PH), which identifies the earliest developmental stage at which future cellular fate becomes significantly predictable beyond random expectation.

Using synthetic systems, lineage-resolved developmental datasets and single-cell multiomic measurements, we demonstrate that prediction horizons vary substantially across biological systems and cellular states. ^13^

Multipotent progenitors exhibit short prediction horizons and high fate entropy, whereas committed cellular states maintain long prediction horizons and low uncertainty.^14^ Furthermore, we show that the addition of increasingly rich molecular information does not uniformly eliminate uncertainty, suggesting that different biological systems possess distinct limits of fate predictability.^15,16^ By quantifying the information-theoretic limits of cell fate prediction, FateLimit establishes a framework for measuring cellular determinism and stochasticity in single-cell systems.^17,18^ More broadly, our approach shifts the focus of single-cell analysis from asking how accurately cell fate can be predicted to asking whether cell fate is fundamentally predictable.

## Results

### FateLimit quantifies the temporal emergence of cellular fate predictability

A central challenge in single-cell biology is determining when future cellular identity becomes predictable from a cell’s current molecular state. Existing trajectory-inference frameworks reconstruct developmental continua and estimate lineage probabilities, yet they do not directly quantify how much information the present cellular state contains about future fate, nor do they identify the developmental stage at which fate becomes statistically predictable. To address this problem, we developed FateLimit, an information-theoretic framework that quantifies the temporal emergence of fate predictability from single-cell omics data (**Fig. 1A–I**). FateLimit operates on a latent representation of cellular state derived from transcriptomic or multimodal measurements and estimates, for every cell, the probability of reaching each terminal fate (**Fig. 1A,B**). Rather than assigning a single deterministic lineage outcome, FateLimit models’ future fate as a probability distribution across all accessible terminal states. From these fate-probability distributions, FateLimit calculates a cell-wise measure of developmental uncertainty termed fate entropy (**Fig. 1C**). Fate entropy is maximal when multiple future outcomes remain equally likely and decreases as cells become progressively committed toward a specific lineage. Consequently, the complement of normalized entropy provides a direct estimate of cell-wise fate predictability. To quantify how information about future fate accumulates during differentiation, we next introduced a mutual-information framework linking current cellular state and future developmental outcome (**Fig. 1D**). For a given developmental interval, fate information was defined as the mutual information between the present state X_t_ and future fate F_t+τ_. This formulation provides a quantitative measure of how strongly future lineage identity is encoded in the current transcriptomic configuration.

**Figure 1.**
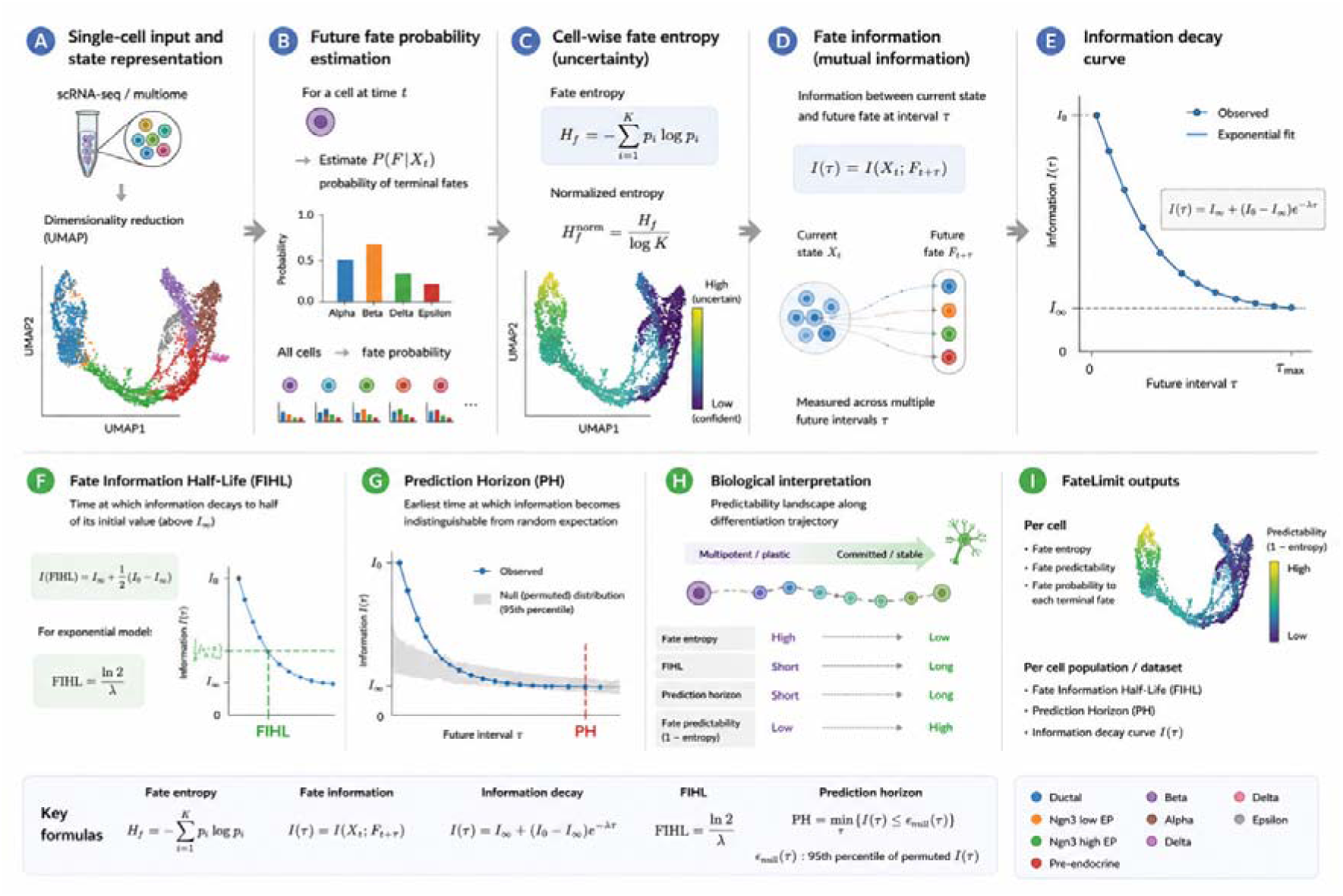
Overview of the FateLimit framework for quantifying the limits of cell fate predictability. (A) Single-cell input and state representation. Single-cell transcriptomic or multiomic measurements are first projected into a low-dimensional latent space representing the current cellular state. This latent representation serves as the starting point for all downstream FateLimit analyses. (B) Future fate probability estimation. For each cell at time t, a probability distribution over future terminal fates is estimated. Rather than assigning a single deterministic outcome, FateLimit models cell fate as a probabilistic distribution across all possible terminal states, yielding. (C) Cell-wise fate entropy. Fate uncertainty is quantified using Shannon entropy of the predicted fate distribution. Cells with multiple competing future outcomes exhibit high fate entropy, whereas lineage-committed cells display low fate entropy. Fate entropy therefore provides a quantitative measure of developmental uncertainty. (D) Fate information. FateLimit measures the mutual information between the current cellular state (X_t_) and future cell fate F_t+T_, I(τ) = I(X_t_;F_t+T_) which quantifies how much information about future fate remains encoded in the present molecular state. (E) Fate-information dynamics. Fate information is evaluated across increasing future intervals τ, generating an information dynamics profile. The dynamics curve characterizes the progressive loss of predictive information during differentiation, development or cellular state transitions. (F) Fate Information Half-Life (FIHL). FIHL measures the characteristic timescale of fate-information dynamics during cellular state transitions. Under an exponential model, FIHL corresponds to the timescale associated with a two-fold change in the dynamic component of fate information. (G) Prediction Horizon (PH) is defined as the earliest developmental stage at which observed fate predictability exceeds the 95th percentile of a permutation-derived null distribution. (H) Biological interpretation. Along differentiation trajectories, multipotent and plastic cell states are expected to exhibit high fate entropy and short prediction horizons. In contrast, committed cellular states display low fate entropy and extended prediction horizons reflecting stable lineage commitment. (I) FateLimit outputs. The framework generates cell-level and population-level measures of cell fate predictability, including fate entropy, fate probabilities, Fate Information Half-Life, Prediction Horizon and information-decay curves. Together, these quantities provide a quantitative description of how rapidly information about future fate is lost from the current cellular state. FateLimit shifts the focus of single-cell analysis from predicting cell fate to quantifying the fundamental limits of cell-fate predictability.

We hypothesized that fate information should change systematically across developmental time. Indeed, theoretical considerations suggest that developmental trajectories transition from highly plastic states with limited predictive information to progressively committed states in which future fate becomes increasingly constrained. To capture this behavior, FateLimit models the temporal dynamics of fate information using an Fate-information dynamics (**Fig. 1E**), allowing estimation of the rate at which uncertainty is resolved during differentiation. Based on this framework, we introduce two quantitative descriptors of developmental commitment. First, the Fate Information Half-Life (FIHL) measures the characteristic timescale over which fate information changes during cellular state transitions (**Fig. 1F**). FIHL provides a system-level estimate of the temporal scale of fate-information dynamics during lineage progression. Second, we define the Prediction Horizon (PH) as the earliest developmental stage at which observed fate information exceeds the 95th percentile of a permutation-derived null distribution (**Fig. 1G**). The prediction horizon therefore represents the point at which future cellular identity becomes statistically distinguishable from random expectation. These metrics enable a unified interpretation of developmental landscapes (**Fig. 1H**). Cells with high entropy exhibit low predictability and short-range information about future fate, consistent with multipotent or plastic states. In contrast, lineage-committed cells display reduced entropy, elevated predictability, and extended prediction horizons. FateLimit thus transforms developmental trajectories into quantitative predictability landscapes, enabling direct comparison of commitment dynamics across biological systems. The framework generates multiple complementary outputs at both the cellular and population levels (**Fig. 1I**). At single-cell resolution, FateLimit estimates fate entropy, fate predictability, and terminal-fate probabilities. At the population level, it provides system-wide measures including the fate information half-life, prediction horizon, and information-decay dynamics. Together, these quantities establish a quantitative framework for measuring when future cellular identity becomes predictable from present molecular state.

### FateLimit reveals the late emergence of fate predictability during pancreatic endocrinogenesis

To demonstrate the ability of FateLimit to quantify developmental predictability in a real biological system, we first applied the framework to a single-cell RNA-sequencing dataset of pancreatic endocrinogenesis comprising ductal cells, endocrine progenitors, pre-endocrine intermediates and terminal endocrine cell types (Alpha, Beta, Delta and Epsilon cells) (**Fig. 2A**). The developmental manifold reconstructed by UMAP recapitulated the established progression from progenitor states toward mature endocrine lineages, providing an ideal system for examining how information about future fate emerges during differentiation. We first quantified cell-wise developmental uncertainty using FateLimit-derived fate entropy. Early progenitor populations exhibited uniformly high entropy values, indicating that multiple endocrine outcomes remained simultaneously accessible from the current transcriptional state (**Fig. 2B**). As cells progressed along the endocrine differentiation trajectory, entropy progressively decreased and became strongly localized around terminal lineage branches. This transition reflects a gradual restriction of developmental potential and suggests that future lineage identity becomes increasingly encoded within the transcriptome during endocrine maturation. To determine whether distinct endocrine fates could be predicted before overt terminal differentiation, we estimated terminal-fate probability distributions for every cell and assigned each cell to its dominant predicted future fate (**Fig. 2C**). Remarkably, lineage-specific territories emerged well before terminal endocrine states became transcriptionally distinct. Cells occupying intermediate regions of the developmental manifold already displayed strong biases toward future Alpha, Beta, Delta or Epsilon identities, indicating that lineage commitment begins substantially earlier than suggested by discrete cell-type annotations alone. We next quantified how developmental predictability changes throughout endocrine differentiation. Cell-wise fate predictability, defined as the complement of normalized fate entropy, increased progressively along latent developmental time (**Fig. 2D**). Early endocrine progenitors exhibited low predictability, consistent with highly plastic developmental states. In contrast, cells approaching terminal endocrine differentiation displayed near-maximal predictability, indicating that future fate became almost completely determined by their current transcriptional configuration. To identify the developmental stage at which future fate first becomes reliably predictable, we calculated the FateLimit prediction horizon. Predictability remained relatively low throughout much of the developmental trajectory before increasing sharply during late endocrine differentiation (**Fig. 2D**).

**Figure 2.**
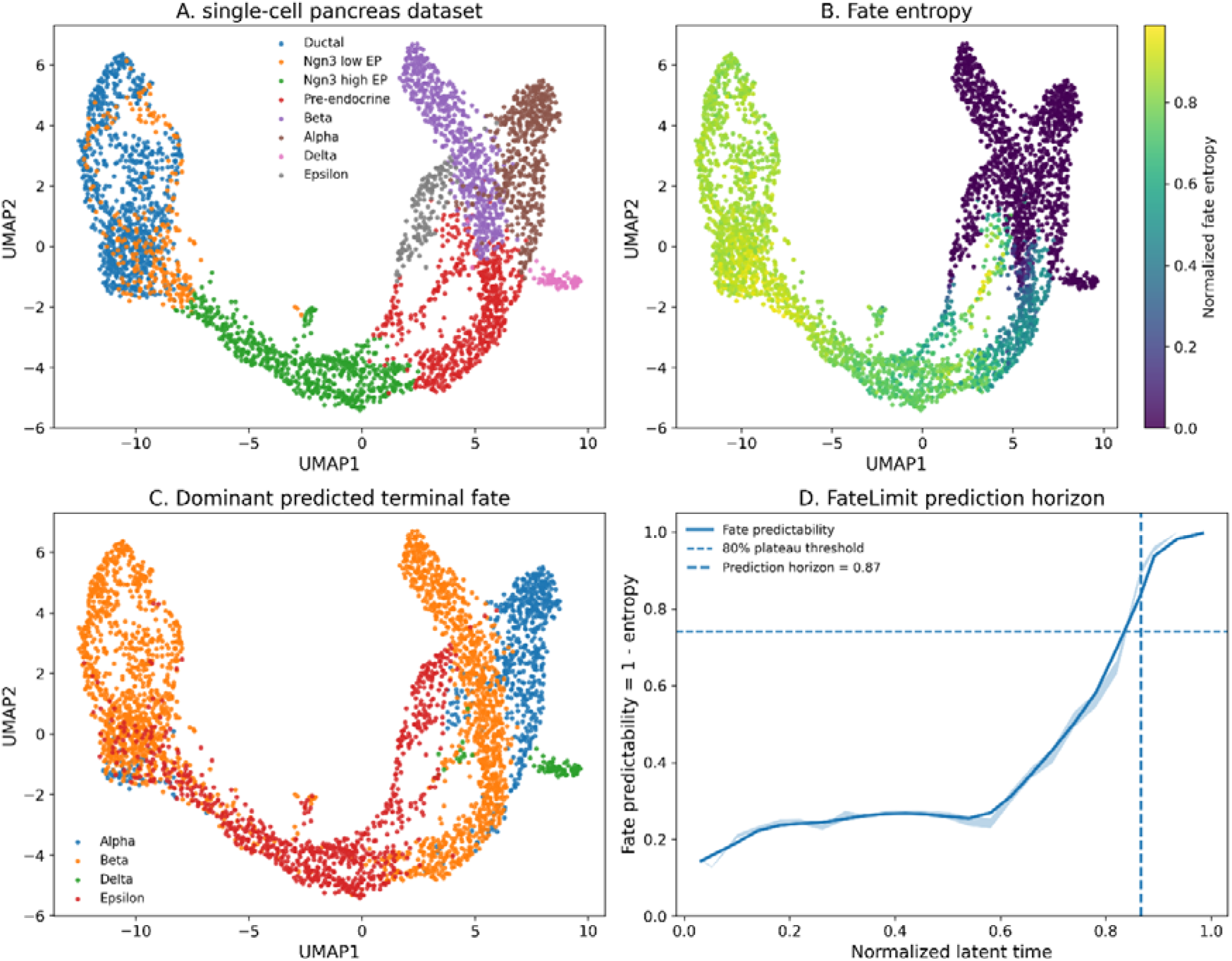
FateLimit quantifies the predictability of cell fate in single-cell transcriptomic data. (A) UMAP visualization of a pancreas endocrinogenesis single-cell RNA sequencing dataset showing major developmental populations, including ductal cells, endocrine progenitors (Ngn3 low EP and Ngn3 high EP), pre-endocrine cells and terminal endocrine cell types (alpha, beta, delta and epsilon cells). The developmental trajectory progresses from progenitor populations toward differentiated endocrine fates. (B) Cell-wise fate entropy projected onto the same UMAP manifold. Fate entropy was calculated from the probability distribution of terminal endocrine fates estimated for each cell. High entropy indicates multiple competing future outcomes and low commitment, whereas low entropy indicates a highly restricted developmental trajectory and increased fate commitment. Early progenitor populations exhibit elevated fate entropy, while terminal endocrine populations display substantially reduced entropy. (C) Dominant predicted terminal fate for each cell. For every cell, terminal fate probabilities were estimated using neighborhood-based assignment in latent transcriptional space. Cells were colored according to the terminal fate with the highest predicted probability. Distinct regions of the developmental manifold progressively acquire lineage-specific fate identities as differentiation proceeds. (D) FateLimit prediction horizon analysis. Fate predictability was defined as one minus normalized fate entropy and was evaluated along latent developmental time. The solid line represents mean fate predictability, and the shaded region indicates variability across cells. The horizontal dashed line denotes the 95th percentile of the permutation-derived null distribution. The vertical dashed line indicates the estimated prediction horizon (0.87 latent-time units), corresponding to the earliest developmental stage at which observed fate predictability significantly exceeds random expectation. These results demonstrate a progressive reduction in developmental uncertainty and an increase in fate commitment during endocrine differentiation.

The prediction horizon was detected at a normalized latent time of 0.87, defined as the earliest point at which observed fate predictability exceeded the 95th percentile of a permutation-derived null model. This result indicates that endocrine lineage identity becomes statistically distinguishable from developmental uncertainty only during the final stages of endocrine commitment.

This observation suggests that pancreatic endocrinogenesis is characterized by prolonged developmental plasticity followed by a relatively abrupt transition toward lineage commitment. Collectively, these results establish pancreatic endocrinogenesis as a system with a late prediction horizon, in which future endocrine identity remains difficult to infer during much of the developmental trajectory despite continuous transcriptional progression. More broadly, the analysis demonstrates that FateLimit converts developmental trajectories into quantitative predictability landscapes and enables direct measurement of when future cellular identity becomes encoded within present molecular state.

### FateLimit quantifies the emergence of fate information during cellular reprogramming

To determine whether FateLimit can quantify predictability in systems characterized by experimentally measured future outcomes, we next analyzed a clonal CellTag-based cellular reprogramming dataset in which cell identities can be linked to their eventual reprogramming fate through lineage tracing (**Fig. 3A**). Unlike developmental trajectory datasets that infer future outcomes computationally, CellTag provides experimentally observed lineage relationships, enabling a direct assessment of whether present transcriptional state contains measurable information about future cellular fate. The reprogramming landscape exhibited substantial transcriptional heterogeneity across sampling time points, reflecting the progressive transition from initial cellular states toward terminal reprogrammed identities (**Fig. 3A**). Cells collected at early reprogramming stages occupied transcriptionally diverse regions of the manifold, whereas later time points became increasingly enriched for cells approaching stable reprogrammed outcomes. Using experimentally observed clonal outcomes as ground-truth future fates, we estimated terminal-fate probabilities for every cell and reconstructed the final reprogramming landscape (**Fig. 3B**). Distinct regions of the transcriptional manifold became associated with alternative future outcomes, indicating that future reprogramming success is partially encoded within present cellular state before terminal conversion is achieved. We next quantified developmental uncertainty using cell-wise fate entropy. Most cells exhibited relatively low entropy values, whereas localized regions of the manifold displayed elevated uncertainty and heterogeneous future outcomes (**Fig. 3C**). These entropy hotspots corresponded to transitional transcriptional states in which individual cells retained access to multiple future reprogramming trajectories. Conversely, cells occupying committed regions of the manifold displayed low entropy and highly predictable outcomes. Because CellTag provides clonal information, we additionally quantified fate diversity at the clone level. Most large clones exhibited near-zero clone entropy, indicating that descendant cells converged toward a common terminal outcome despite transcriptional heterogeneity within the clone (**Fig. 3D**). In contrast, a small number of clones displayed elevated entropy, suggesting incomplete commitment or persistent reprogramming plasticity. These observations demonstrate that fate convergence occurs at the lineage level and provide an orthogonal validation of FateLimit-derived predictability estimates. To investigate how information about future fate emerges during reprogramming, we calculated normalized fate information across successive sampling time points (**Fig. 3E**). Because CellTag provides discrete experimental time points rather than increasing future intervals (τ), we modeled the temporal accumulation of fate information during reprogramming. Fate information increased rapidly during the earliest stages of reprogramming and subsequently reached a stable plateau, indicating that substantial information about future outcome becomes encoded relatively early in the reprogramming process. Exponential fitting revealed an information half-time of approximately 1.36 sampling intervals, suggesting rapid encoding of fate information following reprogramming initiation.

**Figure 3.**
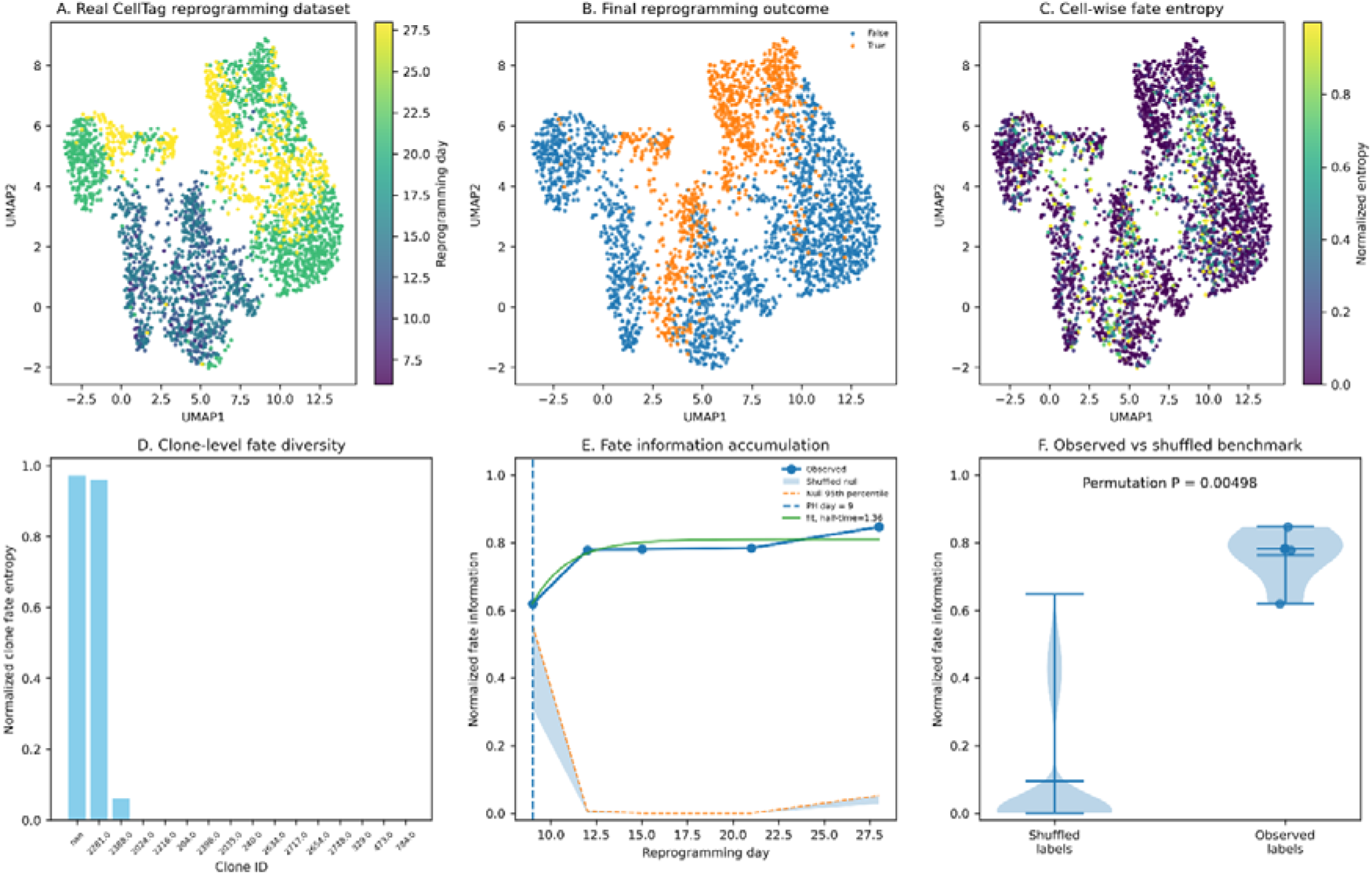
FateLimit detects biologically meaningful fate information in a real lineage-tracing reprogramming dataset. (A) UMAP representation of the Morris–Biddy CellTag reprogramming dataset. Cells are colored according to experimental reprogramming day. Early and late transcriptional states occupy distinct regions of the manifold, illustrating the progressive remodeling of cellular identity during reprogramming. (B) UMAP colored by experimentally observed terminal reprogramming outcome. Cells ultimately classified as successfully reprogrammed or non-reprogrammed occupy partially distinct regions of transcriptional space, indicating that future cellular outcomes become progressively encoded within present molecular states. (C) Cell-wise fate entropy estimated from cross-validated probabilistic prediction of terminal outcome. Fate entropy was calculated as the normalized Shannon entropy of the posterior fate distribution for each cell. Low entropy indicates a highly predictable future outcome, whereas high entropy reflects increased uncertainty regarding future cellular fate. Regions of elevated entropy are concentrated within transitional cellular states, consistent with enhanced fate plasticity during reprogramming. (D) Clone-level fate diversity quantified using CellTag lineage information. For each clone, normalized entropy of terminal outcomes was calculated from the distribution of descendant cell fates. Clones with low entropy predominantly generated a single outcome, whereas high-entropy clones produced heterogeneous descendants, reflecting persistent lineage plasticity and incomplete fate commitment. (E) Fate information accumulation during cellular reprogramming. Normalized mutual information between the current transcriptional state and experimentally observed terminal outcome was quantified at each reprogramming time point. The blue curve represents observed fate information, the shaded region indicates the permutation-derived null distribution, and the dashed orange line denotes the 95th percentile of the null model. Fate information increased progressively throughout reprogramming, indicating that future cellular outcomes become increasingly encoded within the transcriptional state. The vertical dashed line indicates the FateLimit prediction horizon, while the green curve shows the fitted information-accumulation model used to estimate the information half-time. Unlike the FIHL metric used for information-decay analyses, the information half-time in CellTag reflects the rate of information accumulation during reprogramming. (F) Observed-versus-shuffled benchmark. Distribution of normalized fate information obtained using observed outcome labels compared with 200 label-permuted controls. Observed fate information significantly exceeded the null expectation (empirical permutation *P* < 0.005), demonstrating that the measured signal reflects biologically meaningful information regarding future cellular outcomes rather than statistical artifacts arising from dataset structure alone. Collectively, these analyses demonstrate that future reprogramming outcomes are progressively encoded within single-cell transcriptional states. FateLimit quantifies this process through cell-level fate entropy, clone-level fate diversity, information accumulation dynamics, and prediction-horizon analysis, providing a quantitative framework for measuring the limits of cell-fate predictability from single-cell data.

We next evaluated whether the observed fate information exceeded random expectation. A permutation-based null model was generated by randomly shuffling terminal-fate assignments while preserving the overall class distribution. Across all permutations, normalized fate information remained close to zero, whereas the observed data consistently exhibited high information content (**Fig. 3E,F**). The observed value was significantly greater than the shuffled distribution (empirical permutation *P* < 0.005), demonstrating that current transcriptional state contains biologically meaningful information regarding future reprogramming outcome. These analyses establish that FateLimit accurately detects fate information in experimentally validated lineage-tracing datasets and quantitatively measures the emergence of predictability during cellular reprogramming. Unlike developmental systems in which lineage commitment may occur gradually, reprogramming trajectories exhibited rapid information accumulation and early stabilization of fate predictability, highlighting the ability of FateLimit to distinguish distinct modes of cellular state transition.

### FateLimit identifies the emergence of predictability during multilineage hematopoietic differentiation

To evaluate whether FateLimit can quantify predictability in complex developmental systems containing multiple competing lineage outcomes, we next applied the framework to a single-cell RNA-sequencing dataset of human bone marrow hematopoiesis. This system encompasses diverse progenitor and differentiated populations, including erythroid, megakaryocytic, dendritic-cell and myeloid lineages, and therefore provides a stringent test of FateLimit in a highly branched developmental landscape (**Fig. 4A**).

**Figure 4.**
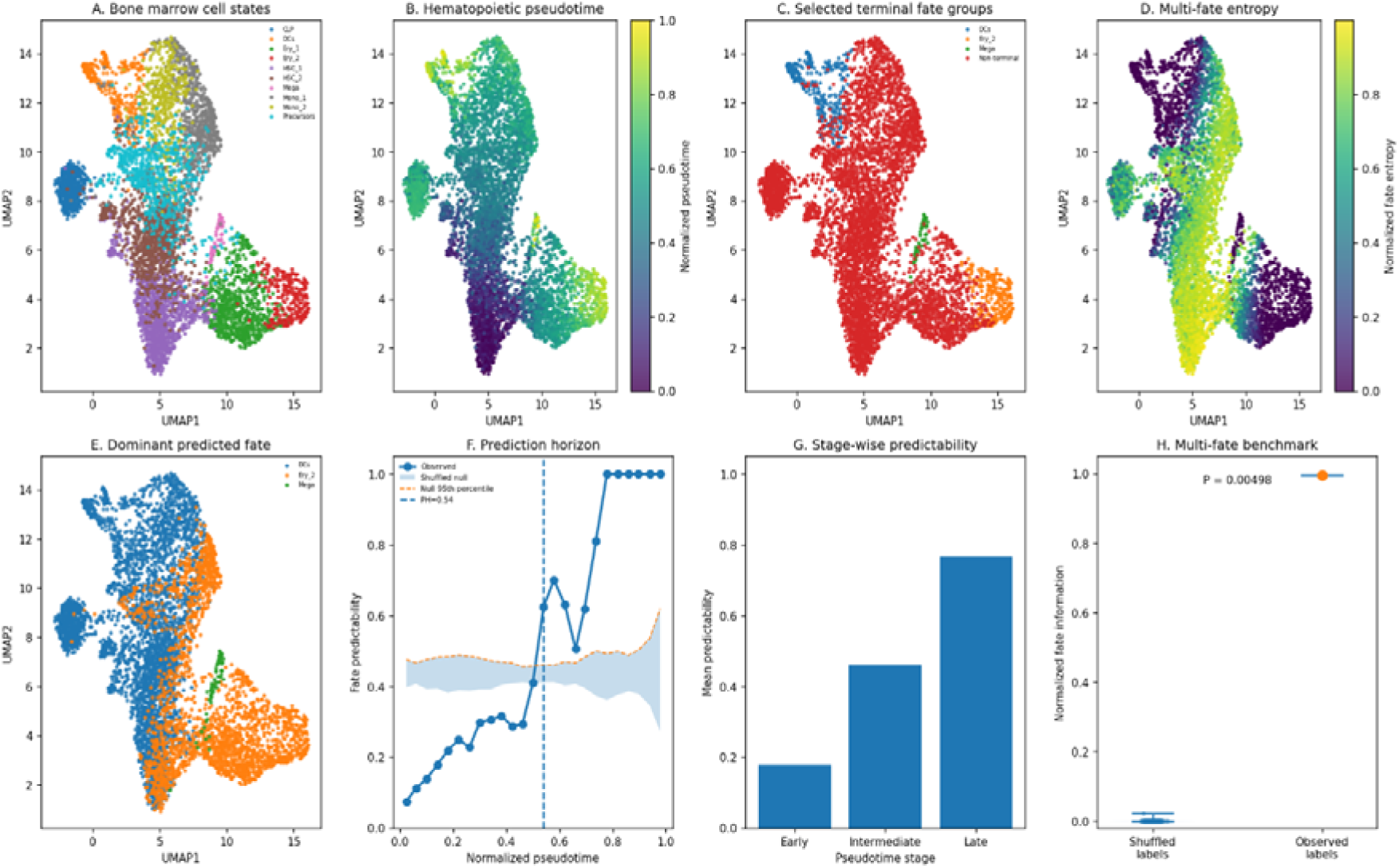
FateLimit quantifies prediction horizons in a multi-lineage hematopoietic differentiation landscape. (A) UMAP representation of a human bone marrow single-cell RNA-sequencing dataset. Cells are colored according to annotated hematopoietic populations, including progenitor, erythroid, dendritic-cell and megakaryocytic lineages. The manifold captures the continuous organization of hematopoietic differentiation and provides a framework for assessing fate predictability across developmental progression. (B) UMAP colored by normalized pseudotime. Pseudotime values were derived from developmental trajectory inference and normalized between 0 and 1. Early progenitor states occupy low-pseudotime regions, whereas differentiated populations are localized at high-pseudotime regions, reflecting progressive lineage commitment. (C) Terminal fate groups used for FateLimit analysis. Late-stage populations enriched at high pseudotime were selected as terminal fates. Cells not assigned to terminal populations were classified as non-terminal states. These terminal populations define the set of competing future outcomes used for probabilistic fate estimation. (D) Multi-fate entropy landscape. For each cell, FateLimit estimated a probability distribution over all terminal fates based on transcriptional similarity to terminal populations. Cell-wise fate entropy was calculated as the normalized Shannon entropy of the resulting fate probability distribution. High-entropy regions correspond to transcriptionally plastic states with multiple competing developmental trajectories, whereas low-entropy regions indicate committed lineage states with highly predictable future outcomes. (E) Dominant predicted terminal fate. Each cell was assigned to the terminal fate with the highest predicted probability. Distinct fate domains emerge across the hematopoietic manifold, demonstrating that transcriptional states contain sufficient information to identify lineage-specific developmental trajectories before full differentiation occurs. (F) Fate predictability across pseudotime. Mean fate predictability was calculated within pseudotime bins and compared with a shuffled-label null model. The solid blue line indicates observed predictability, the shaded region denotes the permutation-derived null distribution, and the dashed orange line marks the 95th percentile of the null model. The vertical dashed line indicates the FateLimit prediction horizon, defined as the earliest developmental stage at which observed predictability significantly exceeds random expectation. Predictability increased progressively along differentiation, indicating that future lineage outcomes become increasingly encoded within transcriptional states. (G) Stage-specific predictability. Mean fate predictability was quantified across early, intermediate and late pseudotime stages. Predictability increased monotonically during differentiation, consistent with progressive restriction of developmental potential and reduction of lineage uncertainty. (H) Multi-lineage benchmark. Normalized fate information obtained from observed terminal-fate labels was compared with shuffled-label controls. Observed fate information substantially exceeded the null expectation (empirical permutation P < 0.005), demonstrating that the inferred fate structure reflects biologically meaningful developmental information rather than stochastic variation. These analyses demonstrate that FateLimit generalizes beyond binary fate systems and can quantify developmental predictability in a complex multi-lineage differentiation landscape. By measuring cell-level fate entropy, dominant lineage bias, and prediction horizons, FateLimit provides a quantitative framework for determining when future cellular identities become predictable during development.

The reconstructed hematopoietic manifold captured the expected progression from precursor populations toward multiple terminal differentiation trajectories (**Fig. 4A,B**). Pseudotime analysis revealed a continuous developmental gradient extending from immature progenitor states to lineage-restricted mature populations, providing a framework for assessing how information about future fate emerges during multilineage differentiation. To define future developmental outcomes, we identified terminal lineage populations corresponding to dendritic-cell (DC), erythroid (Ery_2) and megakaryocytic (Mega) fates and used these populations as reference endpoints for FateLimit analysis (**Fig. 4C**). Unlike binary developmental systems, hematopoiesis presents multiple competing future outcomes, allowing direct assessment of whether FateLimit can resolve lineage-specific predictability in a multifate context. We first quantified developmental uncertainty using cell-wise fate entropy. High entropy values were concentrated within progenitor-rich regions of the developmental manifold, whereas terminal lineage branches displayed markedly reduced entropy (**Fig. 4D**). These results indicate that immature hematopoietic cells retain access to multiple future differentiation trajectories, while lineage commitment progressively restricts the range of accessible cellular outcomes. Notably, entropy decreased continuously along pseudotime rather than abruptly, suggesting a gradual accumulation of fate information throughout hematopoietic differentiation. We next assigned each cell to its dominant predicted future fate based on the maximum terminal-fate probability (**Fig. 4E**). Distinct lineage territories emerged within the developmental manifold, demonstrating that future lineage identity can be inferred prior to terminal differentiation. Cells occupying intermediate regions of the trajectory frequently displayed strong biases toward specific future outcomes, indicating that lineage commitment becomes transcriptionally encoded before acquisition of mature cell-type markers. To quantify when future fate becomes predictable, we calculated FateLimit predictability across pseudotime and compared the observed trajectory to a permutation-derived null model (**Fig. 4F**). Predictability increased steadily during early and intermediate differentiation stages before rising sharply during late pseudotime. The FateLimit prediction horizon was identified at a normalized pseudotime of 0.54, indicating that future hematopoietic lineage identity becomes significantly more predictable than expected by chance approximately midway through the differentiation process. This value is substantially earlier than the prediction horizon observed during pancreatic endocrinogenesis (PH = 0.87), suggesting that lineage commitment emerges earlier in hematopoietic differentiation.

To further characterize the temporal dynamics of commitment, cells were grouped into early, intermediate and late pseudotime intervals. Mean predictability increased monotonically across developmental stages, rising from approximately 0.18 in early progenitors to 0.46 in intermediate states and 0.77 in late differentiation stages (**Fig. 4G**). These results demonstrate progressive accumulation of lineage-specific information as hematopoietic cells approach terminal commitment. Finally, we evaluated whether the observed fate information exceeded random expectation. A permutation benchmark generated by random reassignment of terminal-fate labels produced near-zero information values, whereas the observed data exhibited almost maximal normalized fate information (**Fig. 4H**). The observed value was significantly greater than the shuffled distribution (empirical permutation *P* < 0.005), confirming that current transcriptional state contains substantial information regarding future hematopoietic fate. These analyses demonstrate that FateLimit accurately quantifies fate predictability within a highly branched developmental system and reveals that hematopoietic differentiation is characterized by an intermediate prediction horizon. Unlike pancreatic endocrinogenesis, which exhibits prolonged developmental plasticity before commitment, hematopoiesis accumulates lineage-specific information earlier along the developmental trajectory, highlighting the ability of FateLimit to distinguish distinct modes of fate determination across biological systems.

### Independent validation of FateLimit in vertebrate embryogenesis

To determine whether FateLimit generalizes beyond differentiation and cellular reprogramming systems, we next applied the framework to an independent single-cell RNA-sequencing dataset of zebrafish embryogenesis. This dataset captures a continuous developmental trajectory spanning early embryonic states to multiple differentiated lineages and therefore provides an opportunity to assess whether prediction horizons represent a general property of developmental systems rather than a phenomenon restricted to specific tissues or organisms (**Fig. 5A**). The reconstructed developmental manifold revealed a continuous progression from early embryonic populations toward multiple terminal developmental branches (**Fig. 5A**). Embryonic time annotations were strongly aligned with the inferred developmental trajectory, indicating that the latent manifold accurately captures the temporal progression of vertebrate development. Compared with the pancreatic and hematopoietic systems analyzed above, zebrafish embryogenesis presents a substantially broader developmental landscape encompassing multiple lineage bifurcations and competing terminal outcomes.

**Figure 5.**
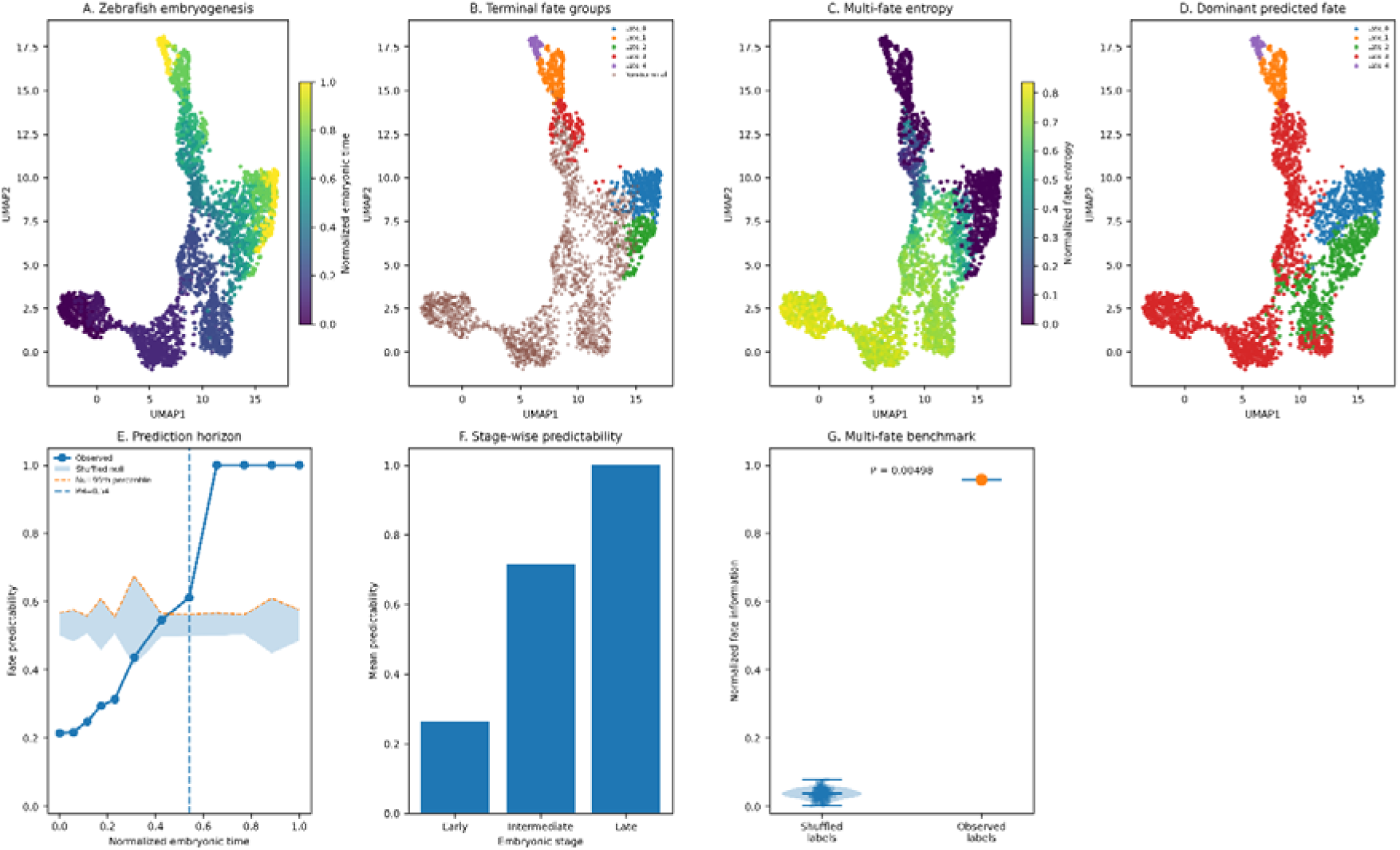
Independent validation of FateLimit in zebrafish embryogenesis reveals the emergence of developmental predictability during vertebrate development. (A) UMAP representation of a real zebrafish embryogenesis single-cell RNA-sequencing dataset colored by normalized developmental time. Cells are organized along a continuous developmental manifold spanning early embryonic states to terminally differentiated populations. The temporal gradient demonstrates progressive transcriptional diversification during vertebrate embryogenesis. (B) Identification of terminal fate groups from late developmental states. Cells located within the latest developmental stages were assigned to terminal fate populations and used as reference outcomes for FateLimit analysis. Cells outside these terminal populations were classified as non-terminal states. These terminal populations define the competing future developmental outcomes used for fate-probability estimation. (C) Cell-wise multi-fate entropy landscape. For each cell, FateLimit estimated a probability distribution across all terminal fates and calculated the normalized Shannon entropy of the resulting distribution. High-entropy regions correspond to transcriptionally plastic progenitor states with multiple competing developmental trajectories, whereas low-entropy regions represent lineage-committed cells with highly predictable future outcomes. Entropy hotspots were concentrated near developmental branching regions. (D) Dominant predicted terminal fate. Each cell was assigned to the terminal fate with the highest predicted probability. Distinct developmental territories emerge across the embryonic manifold, indicating that future lineage outcomes become encoded within transcriptional states before overt terminal differentiation. (E) Fate predictability across embryonic development. Mean fate predictability was calculated along normalized embryonic time and compared with a shuffled-label null model. The shaded region denotes the null distribution, and the dashed orange line indicates the 95th percentile of the null expectation. The vertical dashed line marks the FateLimit prediction horizon (PH = 0.54), defined as the earliest developmental stage at which observed predictability exceeds random expectation. Predictability increased progressively during development, indicating gradual acquisition of lineage-specific information. (F) Stage-wise developmental predictability. Cells were grouped into early, intermediate and late developmental stages based on normalized embryonic time. Mean predictability increased monotonically throughout development, consistent with progressive restriction of developmental potential and increasing commitment toward terminal fates. (G) Multi-fate benchmark against shuffled controls. Normalized fate information derived from observed terminal-fate assignments was compared with a permutation-derived null distribution generated by random label shuffling. Observed fate information was substantially greater than expected by chance (empirical permutation P < 0.005), demonstrating that the inferred developmental structure contains biologically meaningful information regarding future cellular outcomes.

To define future developmental endpoints, we identified five terminal fate groups from late embryonic stages and used these populations as reference outcomes for FateLimit analysis (**Fig. 5B**). These terminal populations represented distinct developmental endpoints emerging during embryogenesis and enabled estimation of future fate probabilities for every cell within the developmental manifold. We first quantified developmental uncertainty using cell-wise fate entropy. High entropy values were concentrated within intermediate developmental regions where multiple future trajectories remained accessible, whereas cells approaching terminal developmental endpoints exhibited markedly reduced entropy (**Fig. 5C**). This pattern is consistent with progressive restriction of developmental potential during embryogenesis and suggests that lineage-specific information accumulates continuously as development proceeds. Importantly, regions of elevated entropy were localized near branching structures of the developmental manifold, supporting the interpretation that these states retain access to multiple competing developmental outcomes. We next assigned each cell to its dominant predicted future fate based on the highest terminal-fate probability (**Fig. 5D**). Distinct developmental territories emerged throughout the manifold, demonstrating that future embryonic outcomes can be inferred from present transcriptional state well before terminal differentiation. Notably, lineage-specific domains became apparent within intermediate developmental regions, indicating that embryonic cells acquire detectable future-fate biases prior to complete lineage commitment.

To quantify the temporal emergence of developmental predictability, we calculated cell-wise fate predictability across normalized embryonic time and compared the observed trajectory to a permutation-derived null model (**Fig. 5E**). Predictability increased progressively throughout development and exceeded the null expectation at a normalized embryonic time of 0.54, defining the FateLimit prediction horizon for zebrafish embryogenesis. Following this transition, predictability rapidly approached maximal values, indicating that future developmental outcomes become strongly encoded within the transcriptional state during the latter half of embryogenesis. We further evaluated developmental commitment by grouping cells into early, intermediate and late embryonic stages. Mean predictability increased monotonically across developmental stages, rising from approximately 0.27 in early embryogenesis to 0.72 in intermediate stages and approaching 1.0 in late developmental states (**Fig. 5F**). These results demonstrate progressive accumulation of developmental information and reveal a continuous transition from highly plastic embryonic states to highly predictable lineage-restricted populations.

To assess statistical significance, we compared the observed fate information to a null distribution generated by random reassignment of terminal-fate labels. The shuffled controls exhibited near-zero information content, whereas the observed data produced almost maximal normalized fate information (**Fig. 5G**). The observed value significantly exceeded the permutation-derived null distribution (empirical permutation *P* < 0.005), demonstrating that the inferred developmental structure contains biologically meaningful information regarding future embryonic outcomes. Finally, we summarized the performance of FateLimit within this independent validation dataset. Analysis of 2,434 cells across 12 developmental time points identified five terminal developmental fates and yielded a prediction horizon of 0.54. The high normalized fate information (0.96) together with significant separation from randomized controls indicates that future embryonic identities become reliably predictable from current transcriptional state during vertebrate development. These results demonstrate that FateLimit generalizes across diverse biological contexts, including endocrine differentiation, cellular reprogramming, hematopoiesis and vertebrate embryogenesis. The observation that zebrafish embryogenesis exhibits a prediction horizon remarkably similar to that observed in hematopoiesis (PH = 0.54) but substantially earlier than pancreatic endocrinogenesis (PH = 0.87) suggests that distinct biological systems possess characteristic timelines of fate commitment. These findings support the concept that prediction horizons represent a general quantitative property of developmental systems and motivate the cross-system comparison performed in Figure 6.

**Figure 6.**
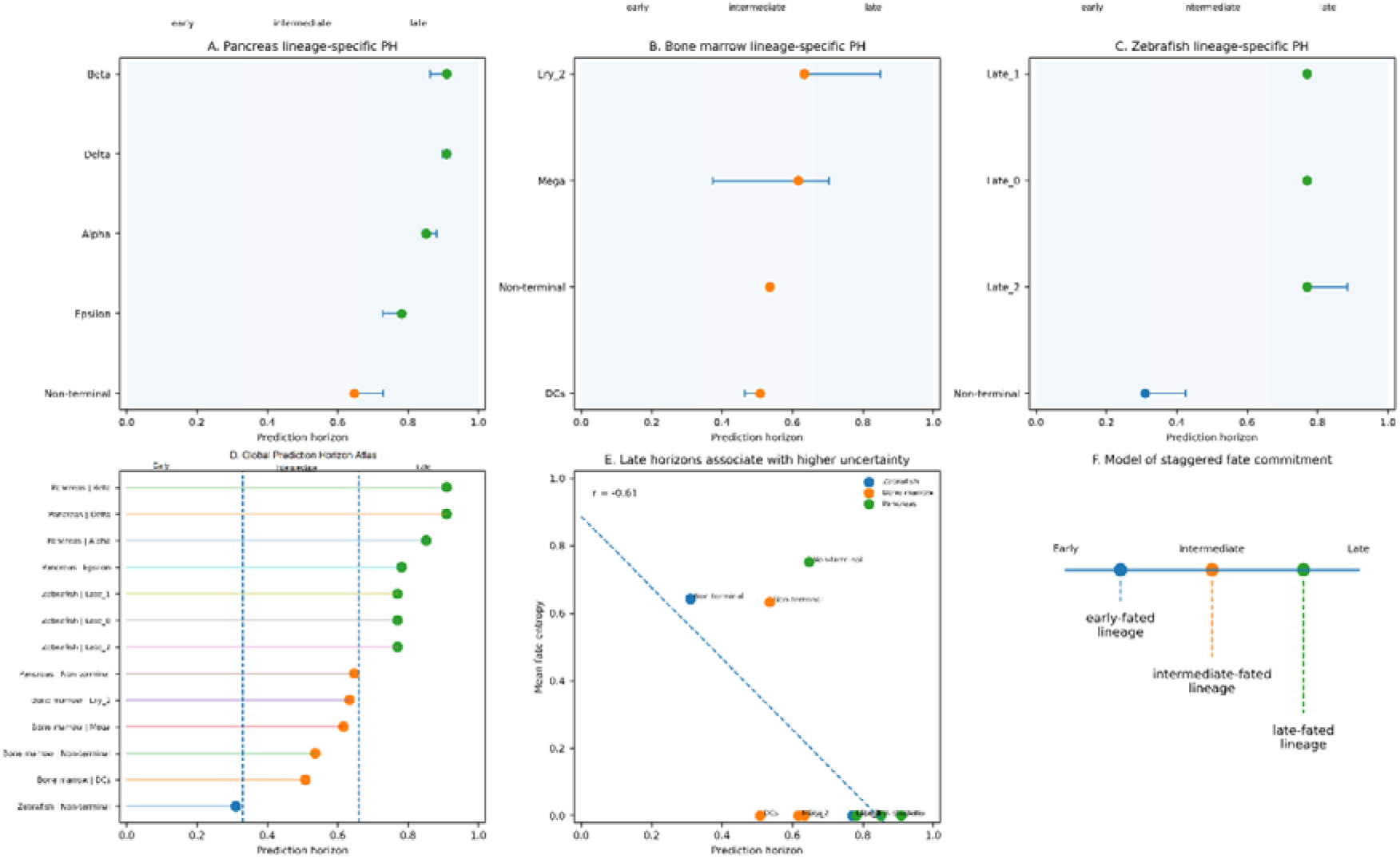
Lineage-specific prediction horizons reveal a hierarchy of fate commitment across biological systems. (A) Lineage-specific prediction horizons in pancreatic endocrinogenesis. For each terminal endocrine lineage (Alpha, Beta, Delta, and Epsilon), a prediction horizon (PH) was estimated as the earliest developmental stage at which future fate became reliably predictable from the current transcriptional state. Points indicate PH estimates and horizontal bars denote bootstrap-derived 95% confidence intervals. Background shading denotes early (0–0.33), intermediate (0.33–0.66), and late (0.66–1.0) developmental intervals. Most endocrine lineages exhibited late prediction horizons, indicating prolonged developmental plasticity before terminal commitment. (B) Lineage-specific prediction horizons in human bone marrow hematopoiesis. Distinct hematopoietic trajectories displayed heterogeneous commitment timing. Dendritic-cell (DC) trajectories became predictable relatively early, whereas erythroid (Ery_2) and megakaryocytic (Mega) lineages exhibited later prediction horizons, suggesting lineage-dependent differences in commitment dynamics. (C) Lineage-specific prediction horizons in zebrafish embryogenesis. Independent embryonic trajectories showed distinct developmental commitment times despite arising within the same organismal context. These results demonstrate that prediction horizons are lineage-specific rather than globally determined by developmental age. (D) Global Prediction Horizon Atlas integrating all analyzed lineages across pancreas, hematopoietic, and embryonic systems. Lineages are ranked according to their prediction horizon values, revealing a continuum from early-committing to late-committing developmental programs. This atlas identifies a hierarchical organization of commitment timing across biological systems. (E) Relationship between lineage-specific prediction horizon and mean fate entropy. Each point represents a lineage-level summary. Lineages with longer developmental uncertainty tended to display altered prediction horizon values, indicating that commitment timing is associated with the persistence of fate ambiguity during differentiation. The dashed line represents the linear regression fit across all analyzed lineages. (F) Conceptual model of staggered fate commitment. FateLimit proposes that cellular differentiation does not proceed through a single universal commitment point. Instead, individual lineages possess characteristic prediction horizons that define when future fate becomes predictable from the transcriptional state. Early-fated lineages become predictable soon after developmental initiation, whereas late-fated lineages retain plasticity over extended developmental intervals before commitment.

### Lineage-specific prediction horizons reveal a hierarchy of fate commitment across biological systems

Having established that FateLimit can quantify developmental predictability in endocrine differentiation, cellular reprogramming, hematopoiesis and vertebrate embryogenesis, we next asked whether different cellular lineages become predictable at similar or distinct developmental times. Although developmental systems are often described using common concepts of commitment and lineage restriction, it remains unknown whether future fate emerges according to a universal temporal program or whether individual lineages possess characteristic commitment dynamics. To address this question, we constructed a cross-system comparison of prediction horizons integrating lineage-specific FateLimit analyses across all datasets examined in this study (**Fig. 6**).

We first estimated lineage-specific prediction horizons for endocrine, hematopoietic and embryonic trajectories. Within pancreatic endocrinogenesis, all major endocrine lineages exhibited remarkably late prediction horizons, with Alpha, Beta and Delta cells becoming predictable only during the final stages of differentiation (**Fig. 6A**). In contrast, hematopoietic lineages displayed substantially earlier commitment dynamics (**Fig. 6B**). Dendritic-cell trajectories became predictable during relatively early developmental stages, whereas erythroid and megakaryocytic lineages exhibited intermediate prediction horizons. Similarly, zebrafish embryonic lineages displayed heterogeneous but generally intermediate commitment timing, with prediction horizons occurring earlier than those observed in pancreatic endocrine differentiation (**Fig. 6C**). To compare commitment timing across biological systems, we ranked all analyzed lineages according to their prediction horizon values (**Fig. 6D**). This analysis revealed a continuous spectrum of developmental commitment rather than discrete categories. Endocrine lineages occupied the extreme late end of the spectrum, whereas several hematopoietic and embryonic trajectories became predictable substantially earlier. These results demonstrate that prediction horizons are lineage-specific properties and suggest that cellular differentiation proceeds through distinct temporal programs of fate commitment. Unexpectedly, prediction horizons varied considerably even among lineages arising within the same developmental system. For example, hematopoietic trajectories exhibited broader variation in prediction horizon than pancreatic endocrine trajectories, indicating that developmental context alone does not determine commitment timing. Instead, individual lineage architectures appear to encode distinct rates of information accumulation during differentiation. To investigate the relationship between developmental uncertainty and commitment timing, we compared lineage-specific prediction horizons with mean fate entropy (**Fig. 6E**). Across all analyzed lineages, prediction horizon exhibited a significant inverse relationship with average entropy (r = –0.61), indicating that lineages characterized by prolonged uncertainty tend to acquire predictive information more gradually. Conversely, lineages exhibiting rapid entropy reduction become predictable earlier during differentiation. These observations suggest that prediction horizons emerge from the balance between developmental plasticity and information accumulation along differentiation trajectories. The resulting framework supports a revised view of cellular commitment (**Fig. 6F**). Classical models frequently assume a single developmental transition separating multipotent and committed states. In contrast, FateLimit reveals that commitment occurs at lineage-specific times and that different cellular identities become predictable according to distinct developmental schedules. Some lineages acquire predictive information early and rapidly restrict their future potential, whereas others retain substantial uncertainty until late stages of differentiation. Collectively, these analyses establish the Prediction Horizon Atlas as a quantitative framework for comparing commitment dynamics across biological systems. More broadly, they reveal a previously unrecognized hierarchy of fate commitment timing in which individual lineages possess characteristic prediction horizons. These findings suggest that the emergence of developmental predictability is itself a measurable biological property and provide a general framework for studying when future cellular identity becomes encoded within present molecular state.

## Discussion

A central goal of single-cell biology is to understand how future cellular identities emerge from present molecular states.^5^ Recent advances in single-cell transcriptomics, trajectory inference, lineage tracing and RNA velocity have enabled increasingly detailed reconstruction of developmental processes. ^5,19^ However, despite these advances, a fundamental question has remained unresolved: when does a future cellular fate become predictable from the current state of a cell? ^10^ Here we introduce FateLimit, an information-theoretic framework that quantifies the emergence of developmental predictability from single-cell omics data. Unlike existing approaches that primarily focus on reconstructing developmental trajectories or estimating lineage probabilities, FateLimit explicitly measures the amount of information that current cellular state contains about future fate and identifies the developmental stage at which fate becomes statistically predictable.^8^ This enables differentiation landscapes to be interpreted not only in terms of cellular state transitions but also in terms of the temporal dynamics of fate information.^20^ A key conceptual advance of FateLimit is the introduction of the prediction horizon, a quantitative measure defining the earliest point at which future cellular identity becomes distinguishable from random expectation. In developmental biology, commitment is often treated as a discrete or qualitatively defined event.^21^ However, commitment boundaries are frequently difficult to identify and may vary depending on the molecular features being examined.^22^ FateLimit reframes this problem as one of information emergence. Rather than asking whether a cell is committed, the framework asks whether sufficient information exists within the current molecular state to predict a future outcome. This perspective provides a quantitative and statistically testable definition of developmental commitment. ^23,24^ Application of FateLimit to pancreatic endocrinogenesis revealed a late prediction horizon, indicating that endocrine progenitors retain substantial developmental uncertainty until the final stages of differentiation.^24–26^ In contrast, hematopoietic differentiation exhibited an earlier prediction horizon, suggesting more rapid accumulation of lineage-specific information.^27–29^ Similar dynamics were observed in vertebrate embryogenesis, where future developmental outcomes became predictable well before terminal differentiation.^22,30^ These findings demonstrate that prediction horizons are measurable properties of developmental systems and are not restricted to a particular tissue, organism or experimental platform.

Importantly, FateLimit also generalized to lineage-tracing data generated by CellTag reprogramming experiments.^31^ In this context, future outcomes were experimentally observed rather than computationally inferred, providing an independent validation of the framework. The ability of FateLimit to recover significant fate information from clonal lineage-tracing data suggests that the framework captures biologically meaningful signals rather than artifacts of trajectory reconstruction.^13^ Together, these analyses indicate that developmental predictability is a robust and quantifiable feature of cellular state transitions. Beyond validating the method across diverse systems, our analyses uncovered an unexpected biological principle. Prediction horizons varied substantially among cellular lineages, even within the same developmental context. Endocrine lineages exhibited consistently late prediction horizons, whereas hematopoietic and embryonic lineages became predictable considerably earlier. The resulting Prediction Horizon Atlas revealed a hierarchy of commitment timing across biological systems. These observations suggest that cellular differentiation does not proceed through a universal commitment point. Instead, individual lineages appear to possess characteristic schedules of information accumulation and fate determination. This concept has important implications for developmental biology. Traditional models frequently describe differentiation as a transition from multipotent to committed states. Our results suggest that commitment may be better understood as a gradual process of information accumulation. Under this view, prediction horizons define the stage at which sufficient information has accumulated to make future fate statistically predictable. Such a framework naturally accommodates continuous developmental trajectories and may provide a quantitative bridge between lineage commitment, transcriptional plasticity and cell-state stability.

Several limitations should be considered. First, FateLimit relies on accurate identification of terminal states and therefore may be sensitive to incomplete sampling of developmental endpoints. Second, prediction horizons are estimated from observed molecular states and do not necessarily imply causal commitment mechanisms. Third, the current implementation primarily uses transcriptomic information and does not explicitly model regulatory, epigenetic or spatial determinants of cell fate. Future extensions incorporating multi-omic measurements, chromatin accessibility, lineage tracing and spatial context may further improve prediction accuracy and biological interpretation. The framework also creates opportunities for applications beyond developmental biology. In regenerative medicine, prediction horizons could identify the optimal stage for directing cell-fate engineering or reprogramming interventions. In cancer biology, prediction horizons may quantify when tumor cells acquire stable malignant phenotypes or treatment-resistant states. More broadly, FateLimit provides a general strategy for measuring predictability in complex biological systems and may be applicable to any process involving transitions between discrete cellular outcomes. In summary, FateLimit introduces a quantitative framework for measuring when future cellular identity becomes predictable from present molecular state. By integrating fate probabilities, entropy, mutual information and prediction horizons, the method transforms developmental trajectories into predictability landscapes. Across differentiation, reprogramming, hematopoiesis and embryogenesis, FateLimit consistently identified measurable prediction horizons and revealed lineage-specific hierarchies of commitment timing. In developmental datasets, FateLimit quantifies the dynamics of predictive information across developmental progression, whereas in CellTag reprogramming data the framework was adapted to quantify the emergence of predictive information across experimentally sampled time points. These findings suggest that the emergence of developmental predictability is itself a fundamental biological property and establish prediction horizons as a general framework for studying cellular fate determination.

## Methods

### FateLimit overview

FateLimit is an information-theoretic framework for quantifying the predictability of cell fate from single-cell measurements. The central aim of FateLimit is not to improve cell-fate prediction accuracy, but to estimate how much information about future cellular outcomes is encoded in the present molecular state and how fate-related information changes across developmental progression over developmental, differentiation or perturbation time. Given a population of single cells measured at time t, each cell is represented by a molecular state vector X_t_. Future cellular outcome is denoted by F_t+τ_, where τ represents a future time interval or developmental distance.^13^ FateLimit estimates the dependence between X_t_ and F_t+τ_ across increasing values of τ. This produces an fate-information dynamics profile from which we derive three quantities: fate entropy, Fate Information Half-Life and Prediction Horizon. The conceptual distinction between FateLimit and existing fate-mapping approaches is that FateLimit asks how predictable cell fate is, rather than which fate each cell is most likely to adopt. Existing approaches estimate transition probabilities, fate probabilities or future transcriptional directions. FateLimit instead quantifies the dynamics of predictive information.

### Input data

FateLimit is applicable to a wide range of single-cell datasets that enable the linkage of current molecular states to future cellular outcomes.^32^ These include longitudinal scRNA-seq datasets, single-cell multiome measurements,^33^ RNA velocity datasets,^5^ clonal lineage-tracing experiments,^34^ CellTag- and LARRY-based lineage-recording systems, perturbation-response datasets with defined terminal phenotypes, and cancer drug-response datasets with sensitive or resistant end states.^35^ The primary requirement is the availability of a cellular state representation and a corresponding future fate annotation that can be assigned at a later time point or developmental stage. For each cell i, the required inputs are: X_i_ the molecular state vector of cell i, t_i_ the sampling time or latent developmental time of cell i, and F_i_ the observed or inferred future fate label. When direct lineage tracing is available, F_i_ is assigned from experimentally observed descendant identity. When direct lineage tracing is not available, F_i_ may be inferred from terminal state assignment or probabilistic fate mapping. In all analyses, direct lineage-tracing labels are preferred over inferred labels.

### Preprocessing of single-cell data

Raw count matrices were filtered to remove low-quality cells and genes.^36,37^ Cells with abnormally low transcript counts, high mitochondrial transcript fractions or extreme library sizes were excluded.^38,39^ Genes detected in fewer than a minimum number of cells were removed. For scRNA-seq data, counts were normalized by total library size, log-transformed and scaled.^40^ Highly variable genes were selected using standard variance-stabilizing procedures. Dimensionality reduction was performed using principal component analysis. Unless otherwise stated, the first 30 principal components were used as the default representation. For multiomic datasets, each modality was preprocessed independently before integration.^41^ Transcriptomic features were processed as described above. Chromatin accessibility features were transformed using term frequency-inverse document frequency normalization followed by latent semantic indexing. ^42,43^ Protein abundance data were centered and scaled. Integrated latent representations were generated using weighted nearest-neighbor integration or variational latent embeddings.^44^ All downstream FateLimit calculations were performed in a latent state space rather than the original high-dimensional gene-expression space, unless otherwise stated. ^45^

### Representation of cellular state

The molecular state of a cell at time t is represented as X_t_ ∈ R^d^ where d is the dimensionality of the latent feature space. FateLimit operates on a generic latent representation of cellular state and is therefore agnostic to the specific feature space used to describe individual cells. Depending on the dataset, X_t_ was defined using principal component coordinates, variational latent embeddings, integrated multiomic representations, ^46^ diffusion map coordinates, ^47^ pretrained single-cell foundation model embeddings, ^48^ or reaction- and pathway-level activity vectors. This formulation enables FateLimit to be applied across diverse single-cell modalities while preserving a common information-theoretic interpretation of cellular state. ^49^ For robustness, FateLimit can be run over multiple representations. ^8^ If multiple representations are available, FateLimit estimates information decay for each representation separately and compares the resulting Fate Information Half-Life values.

### Definition of future fate

Future fate was encoded as a categorical variable F_t+τ_ ∈ {1,2,…,K} where K represents the number of terminal or experimentally defined outcomes. ^6^ Depending on the dataset, fate categories corresponded to endocrine identities (alpha, beta, delta and epsilon cells), hematopoietic lineages (myeloid, erythroid and lymphoid cells), neuronal and glial fates, drug-response phenotypes (drug-sensitive versus drug-resistant states), or metastatic potential (metastatic versus non-metastatic outcomes). For lineage-tracing datasets, fate labels were determined from the observed terminal identities of clonally related descendant cells. ^50^ If a progenitor clone produced multiple terminal fates, the clone was represented by a fate distribution rather than a single fate label. ^51^ For non-lineage datasets, terminal fates were defined using annotated terminal clusters. Fate labels for intermediate cells were inferred using nearest-terminal-cell assignment, Markov absorption probabilities or optimal transport-based mapping. ^52^ These inferred labels were used only for exploratory analysis and were not considered equivalent to experimentally observed lineage fate. ^53^

### Cell-wise fate probability

For each cell, FateLimit estimates a probability distribution over future fate:

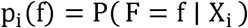

where p_i_(f) is the probability that cell i gives rise to fate f. When lineage tracing is available, this probability is estimated from descendant composition: ^51,54^

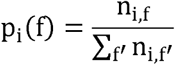

where n_i,f_ is the number of descendants of cell i or clone i assigned to fate f. When only terminal annotations are available, p_i_(f) is estimated using local neighborhood similarity to terminal cells:

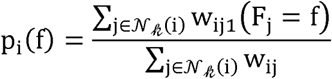

where 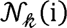 denotes the k-nearest terminal cells of cell i, and ^55^

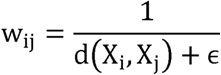

is an inverse-distance weight.

### Fate entropy

Cell-wise fate uncertainty was quantified using Shannon entropy:

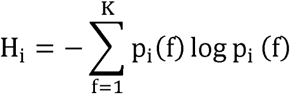

Entropy was normalized by the maximum possible entropy:

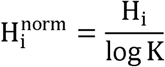

where *K* is the number of possible fates. Thus,

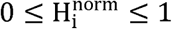

A value close to 0 indicates that a cell has a highly predictable future outcome. A value close to 1 indicates that the cell has multiple competing possible future outcomes. ^14^

Cell-wise fate predictability was defined as: 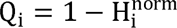 where Q_i_=1 indicates complete predictability and Q_i_=0 indicates maximal fate uncertainty.

### Fate information

To quantify the amount of information about future fate encoded in the current cell state, FateLimit defines fate information as the mutual information between current molecular state and future fate: I(τ) = I(X_t_;F_t+τ_) where X_t_ is the current cellular state and F_t+τ_is the fate observed after temporal interval τ. For discrete variables, mutual information is given by: ^56^

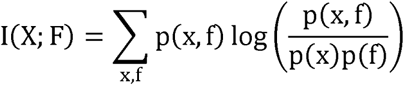

Because single-cell states are continuous, FateLimit estimates mutual information in latent space using either a k-nearest-neighbor estimator or a classifier-based estimator. ^57^

### k-nearest-neighbor mutual information estimator

For continuous X and discrete F, mutual information was estimated using a k-nearest-neighbor estimator. Briefly, for each cell i, the distance to its k-th nearest neighbor in latent space was computed. Local class densities were then estimated within this adaptive neighborhood.

The mutual information was estimated as: ^57^

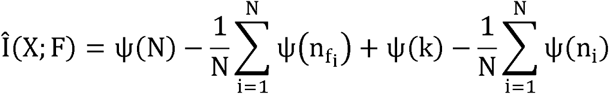

where N is the number of cells, n_fi_is the number of cells belonging to the same fate class as cell i, ni is the number of cells within the local neighborhood of cell i, k is the neighborhood parameter and ψ is the digamma function. Unless otherwise stated, k=20 was used. Sensitivity analyses were performed for k=10,20,30 and 50. ^56,58^

### Classifier-based estimation of fate information

As an alternative estimator, FateLimit uses classifier-derived conditional entropy.

A probabilistic classifier g_θ_ is trained to estimate:

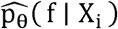

The conditional entropy is then estimated as:

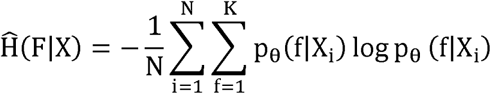

The marginal entropy is:

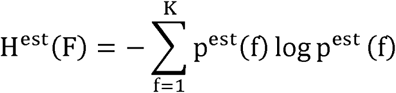

Fate information is then:

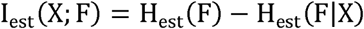

Classifiers used in benchmarking included multinomial logistic regression, random forest, gradient-boosted trees and neural-network classifiers. Model evaluation was performed using cross-validation to avoid optimistic information estimates.

### Fate-information dynamics

FateLimit estimates fate information across increasing temporal intervals: τ_l_,τ_2_,…,τ_m_ For each interval, cells observed at time (t) are paired with future fates at time (t+\tau). This yields an Fate-information dynamics: I(τ_1_), I(τ_2_), …, I(τ_m_) The Fate-information dynamics describes how predictive information changes across developmental progression.^59^

In lineage-tracing data, temporal intervals are defined by experimental sampling time. In static datasets, temporal intervals may be approximated using pseudotime, latent time or experimentally defined differentiation stage. ^9^

### Exponential information-loss model

To summarize information decay, FateLimit fits an exponential model: ^60^

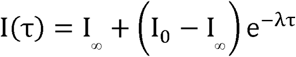

where: I_0_ is initial fate information, I_∞_ is residual long-time information, and λ is the information-loss rate. The model was fitted using nonlinear least squares. Confidence intervals were estimated by bootstrap resampling. ^61^ If I_∞_ was not significantly different from zero, a simplified model was used: I(τ) =I_0_e^-λτ 62^

### Fate Information Half-Life

The Fate Information Half-Life is the time required for fate information to decay to half of its initial value. Using the exponential information-loss model, FIHL is defined as:

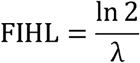

When I_∞_ = 0. For the more general model:

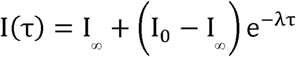

FIHL is defined as the time at which the decaying component reaches half its initial value: ^62^

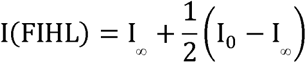

which also gives:

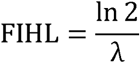

FIHL measures the characteristic timescale over which fate information changes during cellular state transitions.

Interpretation: short FIHL indicates rapid changes in fate information during state transitions; long FIHL indicates gradual changes in fate information across developmental progression; FIHL near zero indicates weak encoding of future fate in the measured state; very long FIHL indicates stable fate restriction and slowly varying developmental information.

### Prediction Horizon

Prediction Horizon (PH) is defined as the earliest developmental stage at which observed fate predictability exceeds the 95th percentile of a permutation-derived null distribution.

Cell-wise fate predictability was defined as Q(t) =1-H_norm_(t), where H_norm_(t) denotes the normalized fate entropy at developmental stage t. To estimate random expectation, fate labels were permuted while preserving the overall cell-state structure. For each permutation (b), fate predictability was recalculated: 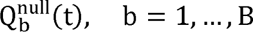 The null threshold was defined as 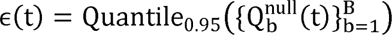 where Q_0.95_ denotes the 95th percentile of the permutation-derived null distribution. Prediction Horizon was then defined as PH = min_t_{t:Q(t) > *ε*(t)} Thus, PH marks the earliest developmental stage at which future cellular fate becomes significantly more predictable than expected by chance. Mutual information was used to quantify the amount of fate-related information encoded in cellular states and to estimate Fate Information Half-Life (FIHL), whereas Prediction Horizon was defined using the emergence of statistically significant fate predictability.^63,64^

### Bootstrap confidence intervals

Uncertainty in FateLimit quantities was estimated using bootstrap resampling. For each dataset, cells or clones were resampled with replacement 1,000 times. For each bootstrap replicate, the full FateLimit pipeline was repeated, including mutual-information estimation, decay-curve fitting, FIHL estimation and Prediction Horizon estimation. The 2.5th and 97.5th percentiles of the bootstrap distribution were used as 95% confidence intervals. When lineage clones were available, resampling was performed at the clone level rather than the cell level to avoid pseudoreplication. ^13,65–67^

### Permutation controls

Permutation controls were used to test whether observed fate information exceeded random expectation. Fate labels were shuffled across cells or clones while preserving: 1. the number of cells; 2. the distribution of cell states; 3. the class frequencies of terminal fates; 4. the sampling-time structure. For each permutation, Fate Information, FIHL and Prediction Horizon were recomputed. Observed values were considered significant if they exceeded the 95th percentile of the permutation distribution. ^57,65^

### Sensitivity analyses

FateLimit estimates were evaluated for robustness to: 1. number of nearest neighbors; 2. latent dimensionality; 3. cell-state representation; 4. terminal fate definition; 5. pseudotime versus experimental time; 6. bootstrap sample size; 7. choice of mutual-information estimator; 8. classifier architecture; 9. class imbalance among terminal fates. For terminal fate imbalance, class-balanced mutual information and stratified resampling were used.

### Application to pancreas endocrinogenesis

FateLimit was applied to a pancreas endocrinogenesis single-cell RNA velocity dataset. Terminal endocrine fates were defined as: alpha cells; beta cells; delta cells; epsilon cells. Cell states were represented using PCA coordinates after standard preprocessing. Latent developmental time was estimated using RNA velocity. Cell-wise terminal fate probabilities were estimated using nearest-terminal-cell assignment in PCA space. Fate entropy, fate predictability and Prediction Horizon were computed across latent developmental time. This analysis was used as an illustrative real-data example and not as definitive lineage validation.

### Application to lineage-tracing datasets

For lineage-tracing datasets, cells sharing the same barcode were grouped into clones. Terminal fates were assigned from the final observed state of each clone. If a clone produced multiple terminal fates, the clone was represented by a multinomial fate distribution. Clone-level Fate Information was estimated using early-state molecular profiles and final descendant fate distributions. FIHL and Prediction Horizon were then estimated using experimentally sampled time intervals. This analysis was used to evaluate whether FateLimit can detect distinct predictability regimes in multipotent and committed cell populations.

### Software implementation

FateLimit was implemented in Python. Core dependencies included: NumPy; SciPy; pandas; scikit-learn; Scanpy; scVelo; AnnData. The package accepts AnnData objects as input and returns cell-wise fate entropy, group-wise Fate Information Half-Life, Prediction Horizon and bootstrap confidence intervals.

### CellTag reprogramming dataset

To evaluate whether FateLimit can identify biologically meaningful information regarding future cellular outcomes, we analyzed the Morris–Biddy CellTag lineage-tracing dataset. This dataset contains single-cell transcriptomes collected during induced cellular reprogramming together with experimentally measured lineage information obtained through combinatorial CellTag barcoding. The dataset was accessed through the CellRank package (cellrank.datasets.reprogramming_morris) and includes multiple reprogramming time points spanning early and late stages of cell-state transition. Cells were annotated according to experimentally observed terminal reprogramming outcomes. Only cells with valid transcriptomic profiles, experimental time-point information and terminal outcome annotations were retained for downstream analyses.

### Probabilistic prediction of future cell fate

For each experimental time point, a probabilistic classifier was trained to predict experimentally observed terminal reprogramming outcomes from the current transcriptional state. Multinomial logistic regression with balanced class weighting was used: P(F | X_t_) Where X_t_ represents the current transcriptional state, F represents the experimentally observed terminal outcome. To avoid overfitting, predictions were generated using stratified five-fold cross-validation. For each cell, the classifier produced a posterior probability vector p= (p_l_,p_2_,…,p_K_) where K denotes the number of possible terminal outcomes.

### Fate information

To quantify how much information regarding future outcomes is encoded in the present transcriptional state, we calculated mutual information between the current state and terminal outcome:

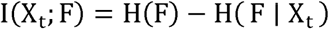

where

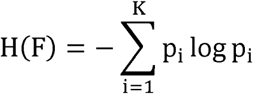

denotes the entropy of the outcome distribution and

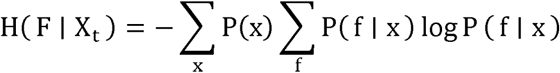

denotes conditional entropy estimated from classifier posterior probabilities. For each experimental time point, normalized fate information was calculated as

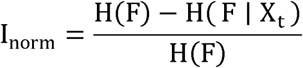

which ranges from 0 (no predictive information) to 1 (perfect predictability).

### Information accumulation dynamics

Unlike developmental trajectory analyses in which fate information is evaluated as a function of increasing future interval (τ), the CellTag dataset consists of discrete experimental sampling time points during reprogramming. Therefore, temporal dynamics were modeled using an information-accumulation framework: I(t) = I∞ − (I∞ − I0)e−λt where I0 is the initial information level, I∞ is the asymptotic information level, and λ is the information-accumulation rate. The information half-time was defined as: t1/2 = ln(2)/λ representing the time required for half of the total attainable fate information to become encoded in the transcriptional state.

### Clone-level fate diversity

To evaluate whether transcriptionally similar cells within a lineage clone generated homogeneous or heterogeneous outcomes, clone-level fate diversity was quantified using CellTag lineage information. For each clone c, terminal outcome entropy was calculated:

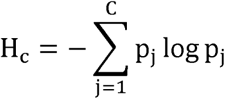

where p_j_ denotes the fraction of clone descendants assigned to outcome j. Entropy values were normalized by the maximum entropy possible for the number of observed outcomes.

Low clone entropy indicates lineage commitment toward a single outcome, whereas high clone entropy indicates persistent fate plasticity within a clone.

### Human bone marrow single-cell dataset

To evaluate whether FateLimit generalizes beyond binary fate systems, we analyzed a human bone marrow single-cell RNA-sequencing dataset representing a continuous hematopoietic differentiation landscape. The dataset was obtained through the CellRank package (‘cellrank.datasets.bone_marrow’) and contains transcriptional profiles spanning progenitor, erythroid, megakaryocytic and dendritic-cell populations together with precomputed developmental pseudotime estimates. Cells lacking valid pseudotime annotations were excluded from downstream analyses. Pseudotime values were normalized between 0 and 1:

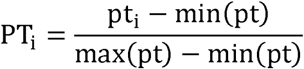

where PT_i_ denotes normalized pseudotime for cell i.

### Identification of terminal fate populations

Unlike the CellTag reprogramming dataset, experimentally observed future outcomes were not available. Therefore, terminal cell populations were identified directly from the developmental manifold. For each annotated cell population, median pseudotime was calculated: M_j_ = median(PT) where j denotes an annotated cell population. Populations enriched at late developmental stages were selected as terminal fates if: M_j_ > Q_0.70_ where Q_0.70_ represents the 70th percentile of median pseudotime across all populations. Populations containing fewer than 50 cells were excluded. The resulting terminal populations constituted the competing future outcomes used for FateLimit analysis.

### Stage-specific predictability

To characterize developmental restriction during differentiation, cells were classified into three pseudotime stages: Early: *PT* < 0.33, Intermediate: 0.33 ≤ PT < 0.66, Late: PT ≥ 0.66 For each stage, mean fate predictability and entropy were calculated. Increasing predictability across pseudotime stages was interpreted as progressive reduction of developmental potential and increasing lineage commitment.

### Zebrafish embryogenesis single-cell dataset

To evaluate the generalizability of FateLimit in an evolutionarily distinct developmental system, we analyzed a publicly available zebrafish embryogenesis single-cell RNA-sequencing dataset distributed through the CellRank framework. The dataset contains transcriptomic profiles spanning multiple embryonic developmental stages and captures continuous transitions from early progenitor states to differentiated cell populations. Cells lacking valid developmental time annotations were excluded. Developmental time values were normalized to the interval [0,1]:

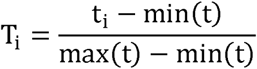

where T_i_ denotes normalized developmental time for cell i.

### Identification of terminal fate populations

Terminal developmental outcomes were identified directly from the developmental manifold. Cells belonging to the latest developmental stages were defined as: T_i_ ≥ Q_0.75_ where Q_0.75_ denotes the 75th percentile of normalized developmental time. Annotated populations enriched within these late developmental states were selected as terminal fate groups. Populations with insufficient representation (<20 cells) were excluded. The resulting terminal populations were treated as candidate developmental endpoints and used as reference fates for FateLimit analysis. For the zebrafish embryogenesis dataset, five terminal fate groups were identified and retained for downstream analyses.

### Stage-specific developmental predictability

To quantify developmental restriction during embryogenesis, cells were divided into three developmental stages: Early: T < 0.33, Intermediate: 0.33 S T < 0.66, Late: Tm 0.66 Mean predictability and entropy were calculated within each stage. Increasing predictability across developmental stages was interpreted as progressive loss of developmental plasticity and increasing commitment toward terminal cell identities.

## Declarations

### Ethics approval and consent to participate

N/A

### Consent for publication

N/A

### Availability of data and materials

The analysis codes and datasets used and/or generated during the current study are available from Dr. Ji-Yong Sung upon reasonable request. Interested researchers may contact Dr. Sung via email at to obtain access to the relevant materials.

### AI Use Declaration

AI tools were used only for English grammar correction and language polishing.

### Authors’ contributions

Conceptualization & Investigation: JYS, JHC; Methodology: JYS; Data analysis: JYS; Writing-original draft: JYS; Writing-review & editing: JYS, JHC; Supervision: JYS, JHC; Project administration: JYS, JHC; Funding acquisition: JHC; Interpretation of the results; JYS, JHC. All authors have read and agreed to the published version of the manuscript.

## Acknowledgements

N/A

